# AFTERHYPERPOLARIZATION AMPLITUDE IN CA1 PYRAMIDAL CELLS OF AGED LONG-EVANS RATS CHARACTERIZED FOR INDIVIDUAL DIFFERENCES

**DOI:** 10.1101/2020.05.15.098376

**Authors:** Daniel Severin, Michela Gallagher, Alfredo Kirkwood

**Author notes:** Correspondence: Michela Gallagher: Department of Psychological and Brain Sciences, 216B Ames Hall, 3400 N. Charles St., Baltimore, MD 21218, Alfredo Kirkwood, Mind Brain Institute, Johns Hopkins University, 250 Dunning Hall, 3400 N. Charles St., Baltimore, MD 21218.

## Abstract

Altered neural excitability is considered a prominent contributing factor to cognitive decline during aging. A clear example is the excess neural activity observed in several temporal lobe structures of cognitively impaired older individuals in rodents and humans. At a cellular level, aging-related changes in mechanisms regulating intrinsic excitability have been well examined in pyramidal cells of the CA1 hippocampal subfield. Studies in the inbred Fisher 344 rat strain document an age-related increase in the slow afterhyperpolarization (AHP) that normally occurs after a burst of action potentials, and serves to reduce subsequent firing. We evaluated the status of the AHP in the outbred Long-Evans rat, a well-established model for studing individual differences in neurocognitive aging. In contrast to the findings reported in the Fisher 344 rats, in the Long-Evan rats we detected a selective reduction in AHP in cognitively impaired aged individuals. We discuss plausible scenarios to account for these differences and also discuss possible implications of these differences.

## INTRODUCTION

Cognitive decline affects a substantial proportion of individuals in the aged population. Research on the neural basis of this decline has revealed that among aged individuals with memory impairments an excess of neural activity is observed in specific circuits within the medial temporal lobe (for review Haberman et al.2017a). Early on, a relation between neural hyperactivity and memory impairment was established from recordings of principle neurons in the CA3 region of the hippocampus in aged Long-Evans rats, where neuronss showed an elevated rate of firing in aged rats with impaired hippocampal-dependent memory while aged cohorts with preserved behavioral performance did not differ from young adults (Wilson et al., 2005b; Wilson et al., 2006). Later studies extended these observations to other mammalian species. Indeed, hippocampal hyperactivity appears to be a dysfunctional condition common to cognitive impairment in aged rodents (Wilson et al., 2005a; El-Hayek et al., 2013), aged rhesus monkeys (Thome et al., 2016), age-related late onset Alzheimer’s disease (AD) in a tau variant mouse model (Maeda et al., 2016), humans with age-related memory impairment (Yassa et al., 2011; Leal and Yassa, 2015) and elderly patients diagnosed with mild cognitive impairment (Yassa et al., 2011).

The manifestation of neural overactivity in age-related cognitive impairment has been localized to hippocampal subregions, especially the dentate gyrus (DG) and CA3 subregions (Haberman et al. 2017b; Simkin et al. 2015; Tran et al. 2019; Wilson et al. 2005a), which receive major input from the layer 2 neurons of entorhinal cortex. At the same time, studies that have shown overactivity in those circuits have not found comparable excess neural activity in the CA1 region, which is innervated by CA3 via Shaffer collaterals (Wilson et al., 2005a; Robitsek et al., 2015). Nonetheless, other studies have reported alterations in the aged rodent CA1. At a synaptic level, animal models with hippocampal hyperactivity and hippocampal-dependent cognitive impairment exhibit alterations in excitatory (Neuman et al., 2015) and inhibitory (Tran et al., 2018) transmission from in vitro studies of CA1. The goal of the current investigation was to examine the intrinsic excitability of CA1 neurons under the modulation of the postburst afterhyperpolarization (AHP), which has been implicated in age-related cognitive impairment.

A context for studies of the AHP in learning was based on the discovery that successful learning in young adult animals was associated with a decrease of the AHP in CA1 pyramidal neurons studied after training in associative eye-blink conditioning (Disterhoft et al., 1986). Many studies, ranging across species and conducted with animals trained in different tasks, subsequently reported an association of decreased AHP with acquisition of learning not limited to the CA1 region of hippocampus (see (Oh and Disterhoft, 2020) for review). This background supported the notion that an alteration in the AHP served as a localized manifestation of learning and a possible contributing mechanism in the acquisition of learning itself. The potential relevance of those findings in the context of aging was initially suggested by an observed augmentation of the AHP in the CA1 region in behaviorally naïve aged animals (Landfield and Pitler, 1984; Disterhoft and Oh, 2007; Kumar and Foster, 2007; Thibault et al., 2007), followed by evidence for age-related differences in AHP as a function of learning performance (Moyer et al., 2000; Tombaugh et al., 2005; Matthews et al., 2009). Such evidence served as a basis for the hypothesis that a greater post-burst AHP in CA1 pyramidal neurons may contribute to a propensity for age-related deficits in learning. While the analysis of AHP in the aged CA1 has been performed primarily in the inbred Fisher 344 rat strain, here it was studied in the outbred Long-Evans strain that has become widely used as a model for individual differences in neurocognitive aging (Barense et al., 2002; Nicholson et al., 2004; Tomas Pereira et al., 2015; Liang et al., 2020).

## METHODS

### Behavioral assessment

Male Long-Evans outbreed rats obtained pathogen-free from Charles River Laboratories (Raleigh, N.C.) were aged 4-6 months (young) or 24-28 months (aged) at the time of behavioral characterization for spatial learning in a water maze (1.83 m diameter, opaque water at 27°C). During an 8-day period, in sessions consisting of 3 trials a day with a 60 seconds intertrial interval, rats were trained to locate a camouflaged platform that remained in the same location 2 cm below the water surface. During a training trial, the rat was placed in the water at the perimeter of the pool and allowed 90 seconds to locate the escape platform. If at 90 seconds, the rat failed to escape on a trial, it was placed onto the platform and allowed to remain there for 30 seconds. The position of entry for the animal was varied at each trial. Every sixth trial consisted of a free swim (“probe trial”), which served to assess the development of a spatially localized search for the escape platform. During probe trials, the rat was allowed to swim a total of 30 seconds with the escape platform retracted to the bottom of the pool. After 30 seconds, the platform was raised so that the rat could complete escape on the trial. A “behavioral index”, which was generated from the proximity of the rat to the escape platform during probe trial performance, was used in correlational analysis with the neurobiological data. This index is the sum of weighted proximity scores measured during probe trials; low scores reflect search near the escape platform whereas high scores reflect search farther away from the target. Thus, the “behavioral index” provides a measure that is based on search accuracy independent of escape velocity (Gallagher and Nicolle, 1993). “Search error” during training trials refers to the deviation from a direct path to the platform and provided an additional measure for behavioral analysis (Gallagher and Nicolle, 1993), Cue training (visible escape platform; 6 trials) occurred on the last day of training to test for sensorimotor and motivational factors independent of spatial learning. Rats that failed to meet a cue criterion of locating the visible platform within an average of 20 seconds over 6 trials were excluded from the experiments.

### Slice preparation

Behavioral groups were intentionally balanced between young, aged unimpaired, and aged impaired rats. The experimenter was blind to the behavioral status of subjects during recording and cellular data analyses. Behaviorally characterized rats were deeply anesthetized with isofluorane, and under Urethane anesthesia (1 g/Kg), they were perfused transcardially with cold dissecting buffer (75 mL at 25 mL/min) containing (in mM) 92 N-methyl-D-glucamine (NMDG), 2.5 KCl, 1.25 NaH_2_PO_4_, 30 NaHCO_3_, 20 HEPES, 25 glucose, 2 thiourea, 5 Na-ascorbate, 3 Na-pyruvate, 0.5 CaCl_2_, and 10 MgSO_4_, pH adjusted to 7.4. After decapitation brains were removed quickly and transferred to ice-cold dissection buffer bubbled with a mixture of 5% CO_2_ and 95% O_2_. The transverse hippocampal slices (300 μm) from the dorsal half of the hippocampus were made using a LeicaVT1200s vibratome. The slices then recovered for 15 min at 30°C and subsequently for 1 hour at room temperature in artificial cerebrospinal fluid (ACSF) in mM: 124 NaCl, 2.5 KCl, 1.25 NaH_2_PO_4_, 26 NaHCO_3_, 25 dextrose, 2 MgCl_2_, and 2.4 mM CaCl_2_, pH 7.4 (bubbled with a mixture of 5% CO_2_ and 95% O_2_). The Institutional Animal Care and Use Committee at Johns Hopkins University approved all procedures.

### Visualized whole-cell voltage-clamp recordings

All recordings were done in a submerged recording chamber superfused with ACSF (34 ± 0.5 °C, 2 mL/min). Visualized whole-cell voltage-clamp recordings were made from CA1 pyramidal cells with glass pipettes (3-5 MΩ) filled with intracellular solution containing the following solutions (in mM): 115 potassium methylsulfate, 20 KCl, 10 Na_2_-phosophocreatine, 10 HEPES, 2 Mg_2_ATP, and 0.3 Na_3_GTP. The junction potential, not corrected, was −6.9 mV. The pH was adjusted with KOH to 7.45; osmolality was between 280 and 290 mOsm. Intrinsic properties were recorded in the presence of 10 μM biccuculine methiodide (Enzo, Life Sciences) and 10 μM saclofen (R&D, Tocris). We studied only cells with series resistance < 25 MΩ, an input resistance > 80 MΩ, resting potential < −58mV, an action potential (AP) height at least 95 mV above the holding potential, and ½ width AP longer than 0.7 ms. Data were filtered at 4 kHz and digitized at 10 kHz using Igor Pro (WaveMetrics Inc., Lake Oswego, OR). All drugs were purchased from Sigma or Thermo Fisher Scientific.

### Statistical analysis

Statistical significance was determined with Prism GraphPad using ANOVA tests followed by Dunnet post hoc test when the data are distributed normally (judged by the D’Agostino-Pearson normality test) or by the Kruskal-Wallis (K-W) test followed by the Dunn’s test.

## RESULTS

The goal of the study was to determine changes in the slow afterhyperpolarization (sAHP) associated with cognitive aging in pyramidal cells of the CA1 hippocampal subfield. To that end we quantified the magnitude of burst-induced sAHP in slices from young mature (6-month old) and aged (24-month old) outbred Long-Evans rats that had been behaviorally characterized in the water maze as described previously (see methods).

The results of the behavioral assessment of the rats used in this study are presented in figure 1A, B. During performance in the first trial (TT data point), before the rats had experienced escaping to the hidden platform in the water maze, the random search to escape was slightly lower in the older rats (p=0.0202 Mann-Whitney test). Subsequently, the young rats were more proficient in learning to locate the platform. A two-way ANOVA (Age x Trial Block) confirmed that performance improved over the course of training (Trial Block, F(3,108) = 34.13, p < 0.0001) and yielded a significant difference between age groups (Age, F(1,108) = 28.26, p < 0.0001). The interaction between Trial Block and Age was not significant, F(3,108) = 1.762, p = 0.159. The learning index scores (Fig. 1B), computed from a key measure of search accuracy during interleaved probe trials, also differed according to age (t(27) = 2.59; p<0.0153) with higher values indicating worse performance. Consistent with previous research in this model, the aged rats displayed a wide spectrum of outcomes, with many aged rats performing on par with young adults and a substantial subgroup performing outside the range of young performance (Fig. 1B). Aged rats performing outside the range of the young group were designated aged impaired (AI), while those performing on par with young adults (Y) were designated aged unimpaired (AU).

**Figure. 1.**
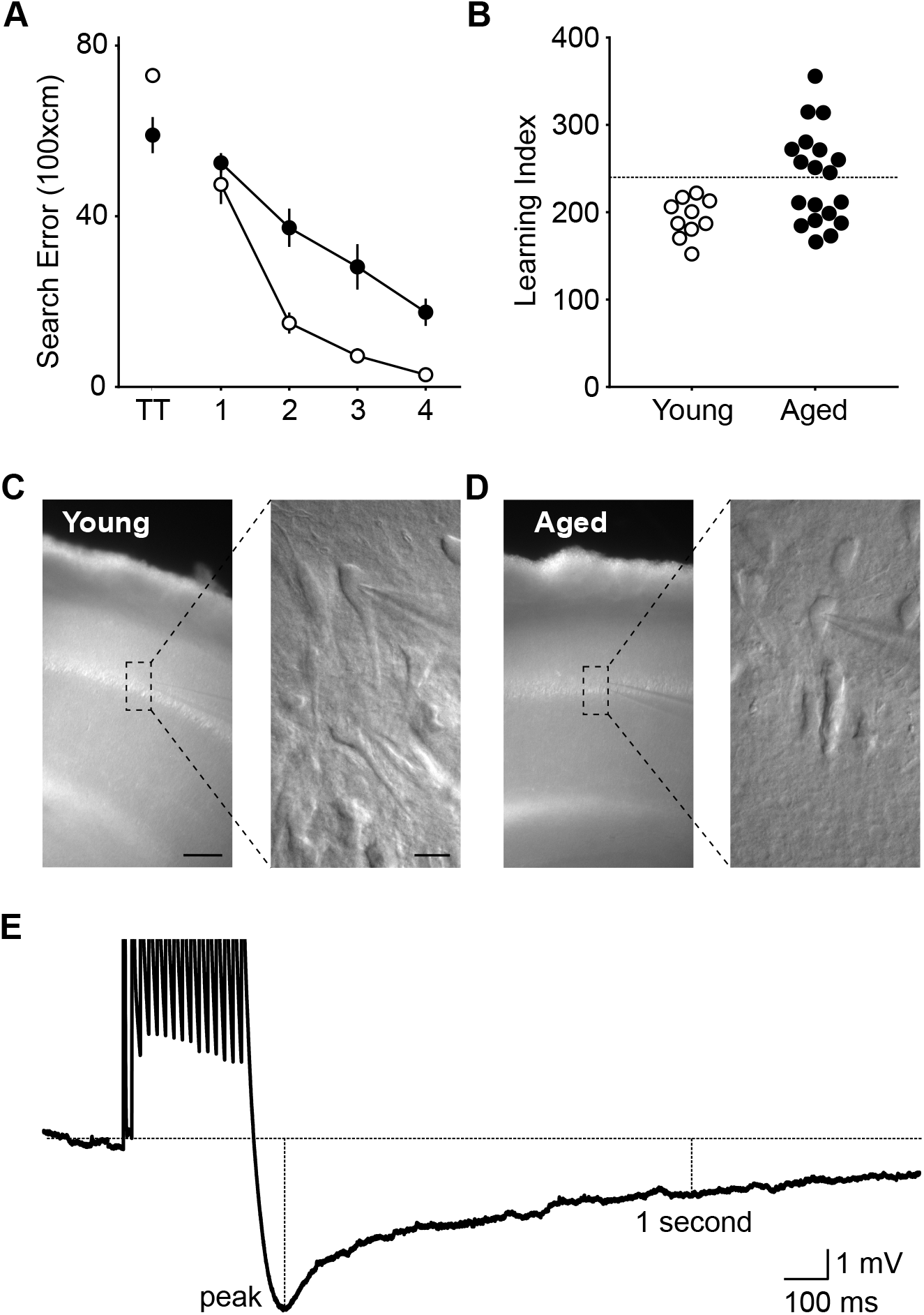
Quantification of sAHP in behavioral characterized young and aged rats. (A) Cumulative search error measure of learning in 5 trial blocks during training in the Morris water maze. This measure reflects the distance of the young (open circles) and aged rats (filled circles) from the escape platform throughout its search, with higher numbers indicating worse performance. Data points represent the average for blocks of 5 training trials ± standard error of mean. Note that during random search prior to the first escape aged rats have proximity scores that are less than young on the first training trial (TT), but young rats are more proficient in learning to escape. (B) A learning index measure for each rat was derived from proximity of the rat’s search during probe trials (see (A), Methods, and Gallagher et al., 1993) with lower scores indicating more accurate performance. As a group, aged rats exhibited significant impairment in accuracy, yet substantial variability. Approximately half of the aged rats performed more poorly than young rats (designated aged impaired), while a substantial subpopulation performed within the range of younger adults (designated aged unimpaired). (C, D) Example pictures of CA1 in slices obtained from a young (C) and an aged rat (D). Images on the right are amplified views of the indicated rectangle. Scale: 100 μm (left), 20 μm (right). (E) Example trace showing the induction of the slow AHP with a burst of action potentials. The dotted lines indicate where the peak and the 1-second measurements were done.

Hippocampal slices were prepared according to previously described procedures for young adult and aged brain tissue (see methods for details) that yield high number of viable cells (Zhao et al., 2011; Tran et al., 2018; Tran et al., 2019) amenable for visualized whole cell recordings (see examples in Fig 1C). To facilitate comparison with previous studies in aged rats with a Fisher background, the sAHP was evoked with the exact same action potential burst protocol (15 APs evoked at 50 Hz with 2 msec suprathreshold current pulses of 1-2 nA) in cells with a targeted membrane potential of −65 mV (see table 1), and in the same ACSF composition as reported previously (Matthews et al., 2009). In each cell recorded we determined the average peak sAHP and the amplitude at 1 sec (see Fig 1D). These averages were computed from at least 20 individual responses to bursts delivered at 0.1 Hz. The results for the young and the aged subgroups (AU and AI) are shown in figure 2. The amplitude of the peak AHP was smaller in the cells of the AI group than in the AU or Y groups (AI: 0.71 ± 0.09mV, n=38 cells, 9 rats; AU: 1.85 ± 0.33mV, n=24,7; 1.59 ± 0.22mV, n=33,10), The significance of these differences was confirmed by a Kruskal-Wallis test (KW stat: 8.64; p= 0.013) and a Dunn’s multiple comparison test (alpha set at 0.05). Similarly, the AHP amplitude at 1 sec after the burst was also affected by aged subgroup (AI: 0.41 ± 0.04 mV, n=38 cells, 9 rats; AU: 0.81 ± 0.14mV, n=24,7; Y: 0.75 ± 0.09mV, n=35,10. KW stat: 7.83, p=0.02), the difference between the AI and the AU and Y groups was confirmed to be significant by the Dunnett’s multiple comparison tests with AI differing from both young and AU, which did not differ from one another. To further examine the differences across the spectrum of aged rat performance, we examined the relationship between each aged individual’s average AHP and its behavioral performance quantified as a learning index (see methods and Figure 1B). We averaged the AHP amplitude (both the peak and amplitude at 1 sec) from all the cells recorded in a given aged individual and plotted this value against the individual’s behavioral performance in the water maze. Only individuals with three or more cells recorded were included in this analysis. As shown in Figure 2C,D the individuals’ averages for both measures of the AHP were significantly correlated with behavioral performance (peak AHP: r^2^ = 0.73, p = 0.0029, Fig 2C; AHP-1sec: t=0.61, p=0.031, Fig 2D), such that higher index behavioral scores indicative of worse performance were associated with more reduced AHP values. In sum, among the aged rats, both measures of the of AHP magnitude are associated with presence/severity of cognitive impairment.

**Table 1.**
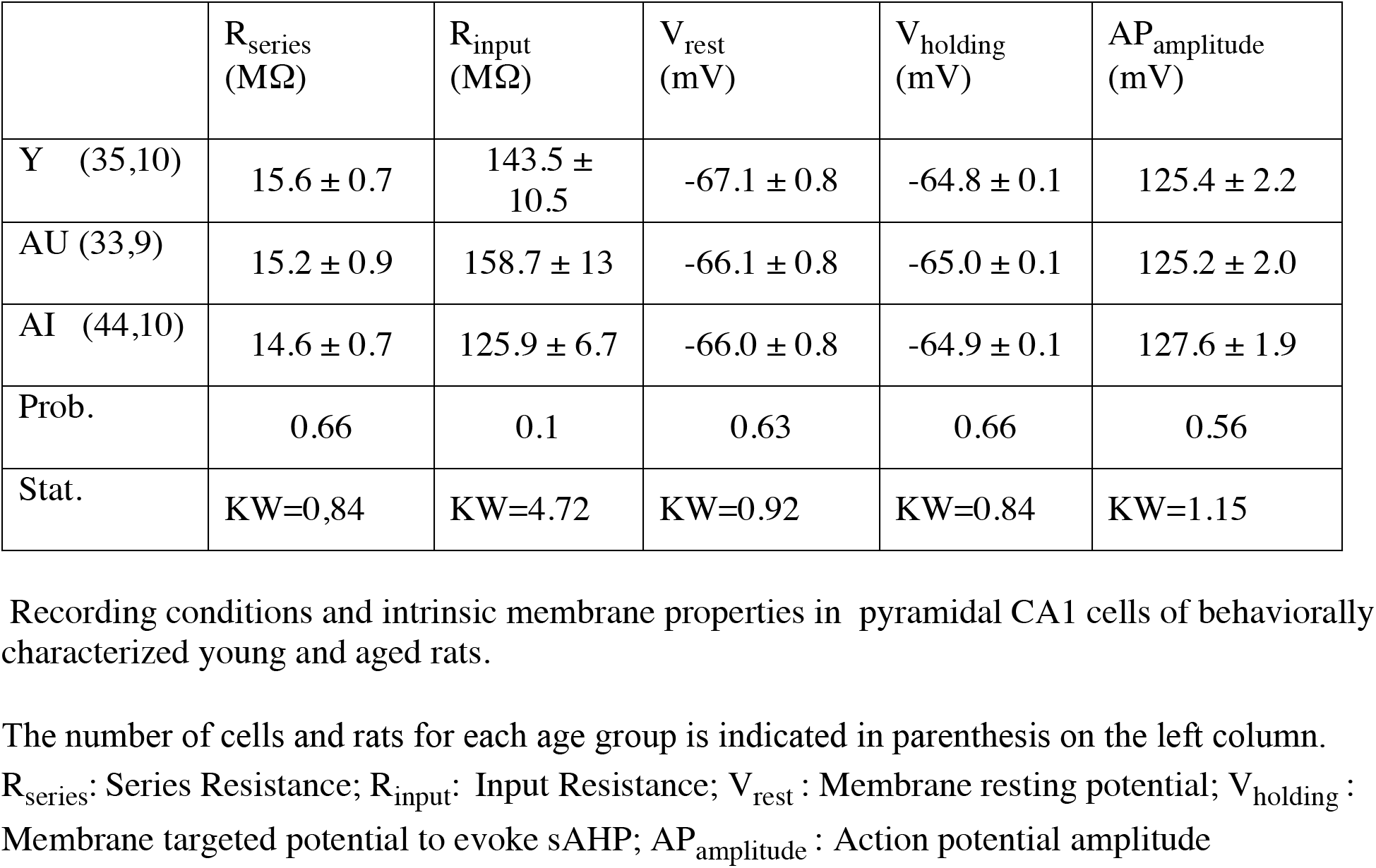
Recording conditions and intrinsic membrane properties in pyramidal CA1 cells of behaviorally characterized young and aged rats.

**Figure. 2.**
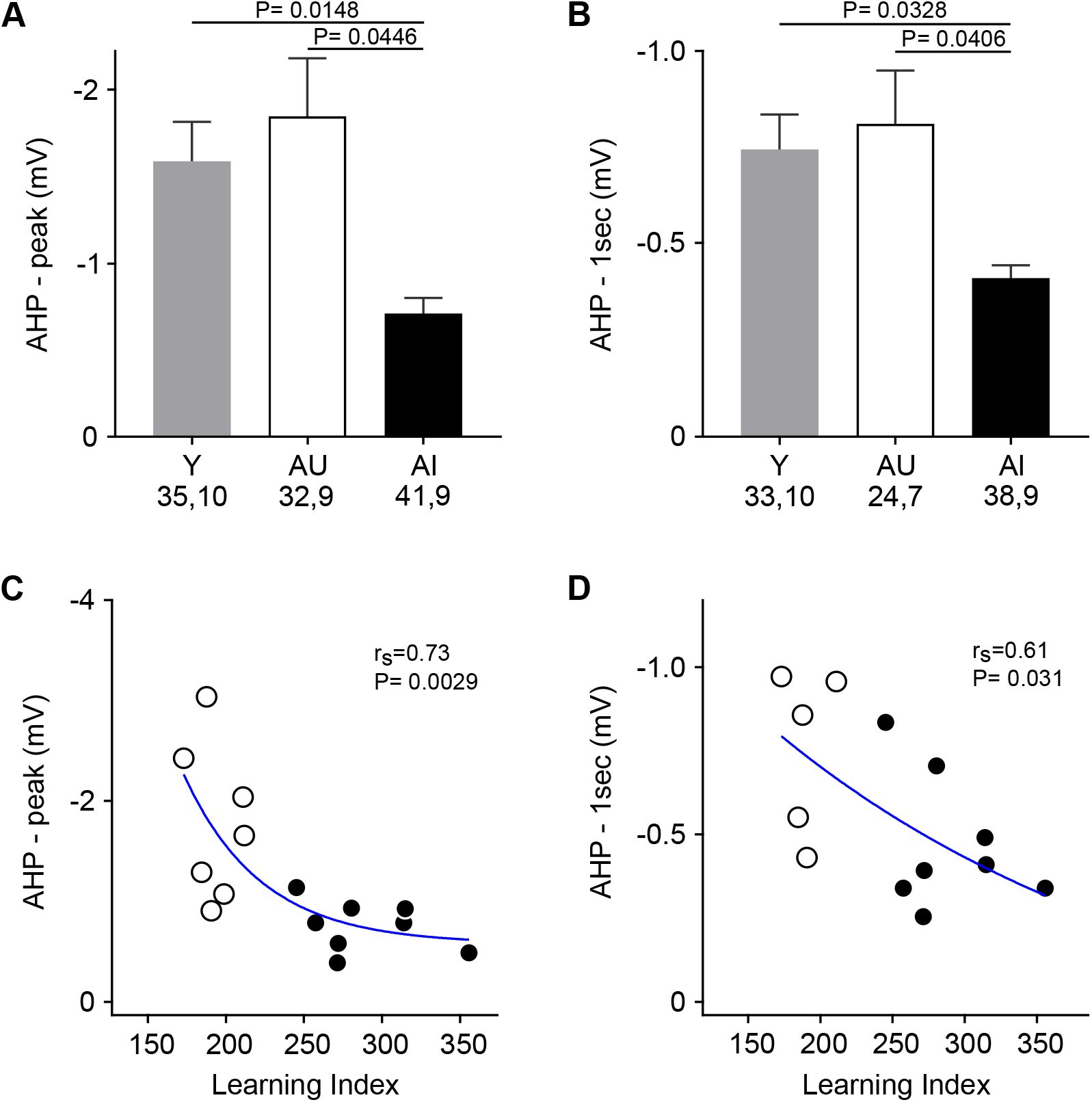
Decreased amplitude of slow AHP in aged impaired animals. Recorded sAHP in CA1 pyramidal cells from young (Y: gray), aged unimpaired (AU: white), and aged impaired (AI: black) rats. Panels on the left (A, C) display AHP – peak data; panels on the right (B, D) show sAHP – 1 sec. (A, B) Average AHP per cell (circles) and average ± standard error of mean of all cells displayed according to each aged subgroup. At the bottom of each column is indicated the number of cells and animals used, and the P value of the comparisons is indicated on the top. (C, D) Relationship between the behavioral performance and the sAHP magnitude averaged across cells in aged individuals (3-9 cells per animal). r_s_ and P, Spearman correlation coefficient and its p–value, respectively. The blue lines in C and D are drawn for visual purposes.

## DISCUSSION

The afterhyperpolarization that follows a burst of action potentials is widely considered to be a feedback mechanism that prevents excess excitability, and it has been extensively studied in CA1 pyramidal cells in the context of learning and aging (Landfield and Pitler, 1984; Disterhoft and Oh, 2007; Kumar and Foster, 2007; Thibault et al., 2007) The present study revealed that in behaviorally characterized outbred aged Long-Evans rats, the AHP magnitude in CA1 is significantly reduced in AI rats, and among all aged rats impairment is significantly correlated with a reduction in measures of the AHP.

The current findings differ from previous studies reporting an age-related increase in AHP magnitude in CA1 cells of aged Fisher 344 rats, with an expected reduction in excitability thought to be a contributing factor to cognitive decline. In one such study the increased AHP was associated with impairment in the water maze performance in the Fisher 344 inbred strain (Tombaugh et al., 2005), which contrasts with the reduction in AHP that we detected in the aged impaired group of outbred Long-Evans rats. The source of this discrepancy is unclear.

Differences in the current results and prior studies might relate to methodological differences in the preparation of the slices. Prior to decapitation we transcardially perfused subjects with solutions developed to optimize visualized whole cell recordings in aged brain tissue, which was not done in the studies on Fisher rats. The interpretation of these discrepant results is further complicated by the fact that the biophysical mechanisms responsible for the sAHP are not fully understood (Pedarzani and Stocker, 2008; Andrade et al., 2012; Gulledge et al., 2013; Oh and Disterhoft, 2020). The studies using aged Fisher 344 rats have emphasized a contribution of Ca-dependent K-channels in the context of the Ca+2-dysregulation hypothesis of aging (Reviewed by (Foster, 2007)(Disterhoft et al., 2004; Thibault et al., 2007). In this view, intracellular Ca+2 accumulates in excess during the AP burst in aged cells resulting in turn in an increased activation of Ca-dependent K-channels to produce a larger AHP. It must be noted, however, that another major contributing mechanism to the AHP, which operates in many neurons of many organisms, including mammalian pyramidal cells, is the pump K^+^Na^+^ATPase responsible for generating and maintaining the Na^+^ and K^+^ transmembrane gradients (Gulledge et al., 2013; Picton et al., 2017; Tiwari et al., 2018). This pump is electrogenic such that an increase on its activity, after a burst of APs for example, produces an outward current that results in a detectable after hyperpolarization (Gulledge et al., 2013). In turn, the pump activity is regulated by multiple signaling pathways (de Lores Arnaiz and Ordieres, 2014) via protein kinases and protein phosphatases (Mohan et al., 2019). In this context, it is important to note that transmembrane ion gradients can be severely disrupted during the preparation of slices. Thus, differences in a cell’s capacity to upregulate the K^+^Na^+^ATPase to restore gradients might, in itself, contribute to the differences in AHP observed across age groups. In a similar fashion, the different methods used to prepare slices in the studies of Long-Evans and Fisher 344 rats might affect ion gradients differently, eliciting different compensatory cell responses, which might have in turn contributed to the different outcomes reported.

An alternative explanation to account for the AHP results in Fisher 344 and Long Evans rats is that the differences relate to different trajectories of aging in the two rat models. Besides changes in AHP, multiple mechanisms participate in controlling the homeostasis and plasticity of intrinsic excitability in neural circuits (for reviews see (Misonou, 2010; Titley et al., 2017; Reuveni and Barkai, 2018; Debanne et al., 2019). During aging the recruitment of these mechanisms in CA1 might differ among rat strains, such that a similar set point of excitability is achieved in different ways. Yet another possibility is that learning requires AHP in CA1 to be within a “permissive window”, above or below which learning is impaired. Notably, the CA1 AHP magnitude of young adult Fisher 344 rats (Matthews et al., 2009) is larger than in Long-Evans (Figure2) rats. Thus, if the AHP magnitude in Fisher rats sits close to an upper tolerance limit, increases in AHP will diminish performance. Conversely, if AHP in Long-Evans rats sits close to a lower tolerance limit, a decrease in AHP would compromise learning.

In this regard, it is worth noting that changes in neural excitability can have differential effects according to age. One case is the pharmacological regulation of tonic GABAergic inhibition in the hippocampus. Reducing tonic inhibition has been shown to have beneficial cognitive effects in young adults but the same treatments are not efficacious in aged animals (Martin et al., 2010)(Koh et al., 2013). Indeed reduction of tonic inhibition in high-performing aged Long Evans rats impairs performance (Koh et al., 2020) and positive allosteric modulators that increase tonic inhibition improve performance in aged rats that are otherwise impaired (Koh et al., 2013). Interestingly, high performing aged Long Evans rats also exhibit an increased tonic inhibition localized to CA1 pyramidal neurons (Tran et al., 2018). As another example, increasing GABAergic inhibition in very immature visual cortex enables experience-dependent plasticity, yet the same manipulation prevents plasticity in mature cortex (Feldman, 2000; Jiang et al., 2005).

In sum, we pose that the different in age-related changes in slow AHP in CA1 reported in Fisher 344 and Long-Evans rats open interesting questions concerning excitability homeostasis during aging, yet the limited understanding of the biophysical mechanisms underlying the slow AHP, preclude generalizing results of any particular strain.

## ACKNOWLEDGMENTS

This work was supported by National Institute on Aging/National Institutes of Health Grants AG009973-22 to MG and AG009973-24 to AK.

## Notes

<u>Acknowledgments</u>:. Supported by grant PO1-AG09973 from NIH.

### Competing Interest Statement

The authors have declared no competing interest.

